# The *Prrx1* limb enhancer marks an adult population of injury-responsive dermal fibroblasts

**DOI:** 10.1101/524124

**Authors:** Joshua D. Currie, Lidia Grosser, Prayag Murawala, Maritta Schuez, Martin Michel, Elly M. Tanaka, Tatiana Sandoval-Guzmán

## Abstract

The heterogeneous properties of dermal cell populations have been posited to contribute toward fibrotic, imperfect wound healing in mammals. Here we characterize an adult population of dermal fibroblasts that maintain an active *Prrx1* enhancer which originally marked mesenchymal limb progenitors. In contrast to their abundance in limb development, postnatal *Prrx1* enhancer-positive cells (Prrx1^enh+^) make up a small subset of adult dermal cells (∼0.2%) and reside mainly within dermal perivascular and hair follicle niches. Lineage tracing of adult Prrx1^enh+^ cells shows that they remain in their niches and in small number over a long period of time. Upon injury however, Prrx1^enh+^ cells readily migrate into the wound bed and amplify on average 16-fold beyond their uninjured numbers. Additionally, following wounding dermal Prrx1^enh+^ cells are found out of their dermal niches and contribute to subcutaneous tissue. Postnatal Prrx1^enh+^ cells are uniquely injury-responsive despite being a meager minority in the adult skin.

## Introduction

The skin is the largest organ and one with a crucial task: making a multifunctional barrier between internal organs and the outside environment. The regenerative capacity of the skin is essential to maintain its integrity. However, in adult skin wound healing typically results in scar tissue. In the search for therapies that enhance wound healing or reduce fibrosis, cellular heterogeneity has emerged as an added layer of complexity that profoundly shapes the outcome of wound responses (Driskell et al., 2013). The heterogeneity of fibroblasts has been of particular importance, since these cells are the main actors during wound healing to produce either scar-free healing or unresolved fibrotic scars (Gurtner et al., 2008). Central to this idea, is that mixed populations of cells may carry intrinsic differences in their response to wound-related signals or their capacity to reconstitute all the structures of the intact organ.

Several groups have identified fibroblast subgroups by a single or battery of molecular markers. Cells derived from Engrailed-1 embryonic lineage (Rinkevich et al., 2015) or adult cells with *Gli-1*^+^ expression have been shown to contribute to wound fibrosis (Kramann et al., 2014), while Wnt-dependent expansion of BLIMP1^+^ dermal cells can support de novo hair follicle formation during wound repair (Kretzschmar et al., 2014). Ideally, such categorization would distinguish subpopulations, with higher regenerative or differentiation potential, that could be analyzed in isolation from fibrosis-associated cells. The final goal would be to amplify and recruit non-fibrotic populations during wound repair, or inversely, deter fibrotic cells from making contributions to wound healing.

To identify adult cells that retain a progenitor-like ability to participate in tissue formation, we looked at molecular markers that are present during organogenesis. One such marker is the transcription factor paired-related homeobox 1 (*Prrx1* or *Prx1*), an early marker of lateral plate mesoderm (LPM) that labels progenitors of nascent limb skeleton and soft connective tissue of the flank and limb. Additionally, *Prrx1* is upregulated following salamander limb amputation (Satoh et al., 2007) as well as in anuran limb regeneration (Suzuki et al., 2005). Transgenic mouse models of *Prrx1* activity have relied on a specific enhancer that encompasses approximately 2.4kb upstream of the transcriptional start site (Logan et al., 2002). In reporter lines, this enhancer was used to drive LacZ or Cre recombinase expression in embryonic lateral soft connective tissue, portions of craniofacial mesenchyme, and limb skeleton and connective tissue. A recent report has implicated a population of PRRX1^+^ cells in the regeneration of calvarial bone (Wilk et al., 2017), but whether PRRX1 protein (PRRX1^+^) or enhancer activity (Prrx1^enh+^) remain postnatally in other tissues is unknown. This led us to investigate *Prrx1* expression and enhancer activity in the skin to determine its role in homeostasis and tissue repair.

## Results and Discussion

### PRRX1 protein marks a broad population of limb bud progenitors and adult mesenchymal dermal cells

*Prrx1* was originally characterized as a progenitor marker of limb skeleton and soft connective tissue using a combination of *in situ* hybridization and Cre activity or LacZ expression in reporter mice (Chesterman et al., 2001). However, a precise timeline of protein expression at both embryonic and postnatal timepoints is unknown. To do this, we used a previously characterized polyclonal antibody anti-PRRX1 (Gerber et al., 2018; Oliveira et al., 2017). By immunohistochemistry, PRRX1**^+^** cells were detected in limb bud and lateral plate at E9.5, where most mesenchymal cells are positive (Figure 1 A). At this stage, expression of the protein is homogenous in what is considered the beginning of the budding phase. At E10.5 the limb bud is defined and protruding from the body flank. PRRX1**-** cells exist at the junction of the arm, possibly from colonizing endothelial and myogenic precursors (Figure 1 B). At E12.5, cartilage condensations become evident, with cells within the condensate (SOX9+ cells) down regulating *Prrx1* expression. However, most mesenchymal cells still remain PRRX1**^+^** (Figure 1 C).

**Figure 1.**
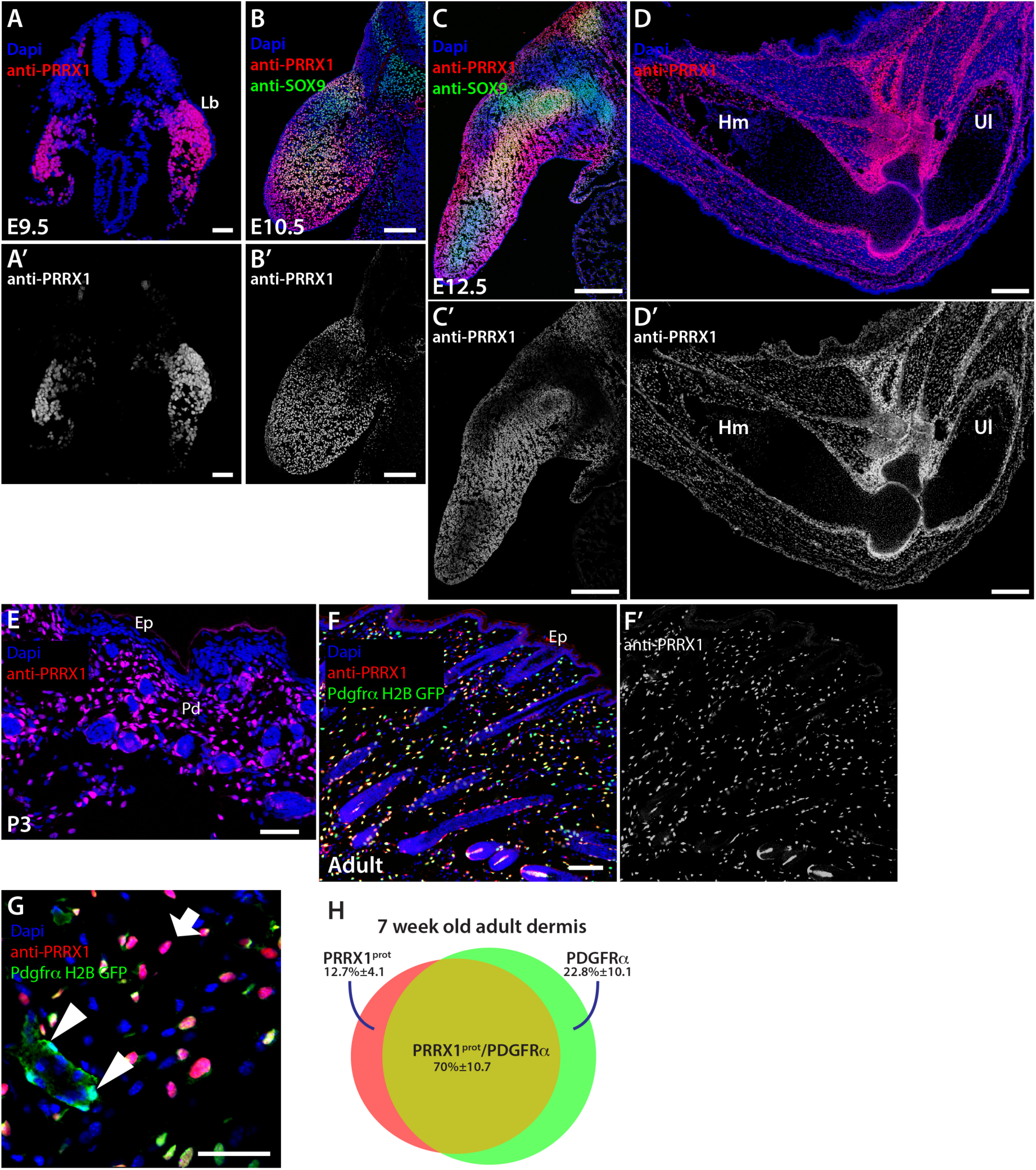
PRRX1 protein marks a broad mesenchymal population during limb development and in adult dermal tissue. (A, A’) Representative micrographs of antibody staining against PRRX1 protein. The peak of PRRX1 in the limb bud (Lb) is around embryonic day E9-10, nuclei in blue, PRRX1 antibody staining in red, greyscale (A’). Scale bar, 50 microns. (B, B’) At E10.5 cartilage condensations, positive for SOX9 protein (in green), in the midline of the limb down regulate PRRX1 protein. Scale bar, 200 microns. (C, C’) By E12.5, skeletal condensations are distributed along the limb and downregulate PRRX1. Scale bar, 500 microns. (D, D’) At E16.5, the limb has patterned the musculo-skeletal elements Humerus (Hm), a clear elbow joint, Ulna (Ul) and digits. PRRX1 is highest at the elbow area. Scale bar, 200 microns. (E) After birth, at postnatal day 3 (P3), PRRX1^+^ cells are still present across dermis, including reticular and papillary dermis (Pd). Epidermis (Ep) is negative for PRRX1. Scale bar, 50 microns. (F) In adult skin, PRRX1^+^ cells in red, greyscale (F’) are compared to the population of cells PDGFRα^+^ in green and quantified (H). Scale bars (F) 200 microns. (G) High magnification of adult skin. Arrows mark PDGFRα^+^ cells that are PRRX1^−^. Arrowheads, PDGFRα^−^ cells that are PRRX1^+^. Scale bar, 50 microns. (H) Quantification of the PDGFRα and PRRX1 populations in adult dermis, represented in a Venn diagram. The mean percentage of cells/mm^2^ ± SD is reported.

At E16.5, clear PRRX1**^+^** and PRRX1**-** zones were visible in the limb, although most connective tissue cells were still PRRX1**^+^** (Figure 1 D). We further investigated if PRRX1 remains in postnatal tissue or if its expression is restricted to embryonic and neonatal stages. In postnatal day 3 (P3) PRRX1**^+^** cells persist abundantly in the dermis (Figure 1 E). Since PDGFRα has been previously suggested as a pan marker of dermal fibroblasts (Driskell et al., 2013), we used the *Pdgfrα-H2B-EGFP* transgenic mouse to quantify the overlap of PRRX1**^+^** in adult mesenchymal dermal tissue (Figure 1 F). We found that in 7 week-old mice 12,7% ± 4,1 (n=6) of total PRRX1**^+^** cells are PDGFRα**^−^**. Conversely, 22,8% ± 10,1 of the total PDGFRα**^+^** cells are PRRX1**^−^** (Figure 1 G-H). This result was consistent with data from Rinkevich et al., who found that a considerable percentage of *Engrailed-1* lineage fibroblasts do not express PDGFRα (Rinkevich et al., 2015). Neither PRRX1 nor PDGFRα encompassed the full complement of mesenchymal cells and PDGFRα is also expressed in non-mesenchymal lineages such as megakaryocytes and platelets (Demoulin and Montano-Almendras, 2012; Ye et al., 2010). Therefore, our data shows that PRRX1 is an optimal marker to demarcate a broad mesenchymal population in the dermis.

### The *Prrx1* enhancer labels embryonic mesenchymal progenitor cells and a small subset of the broad PRRX1 dermal cell population

To trace the fate of PRRX1 cells in homeostasis and injury, we generated transgenic mice expressing Cre-ERT under the control of the 2.4kb *Prrx1* enhancer (Logan et al., 2002) together with nuclear teal fluorescent protein (TFP) as a reporter. We next crossed the *Prrx1 enhancer-CreER-T2A-mTFPnls* mice with *Rosa-CAG-loxP-stop-loxP-TdTomato* mice (referred to as *Prrx1enh-CreER;LSL-tdTomato*)(Figure 2 A), allowing us to trace the fate of *Prrx1* enhancer-positive cells (Prrx1^enh+^) upon tamoxifen administration. A single low dose of tamoxifen (1mg, gavage) to gestating mothers to elicit conversion in E10.5 embryos, yielded labeling of Prrx1^enh+^ cells in a majority of limb bud cells, facial mesenchyme, and inter-limb flank (Figure 2 B). In postnatal and juvenile mice, the conversion of the *Prrx1* enhancer was vastly reduced such that even after repeated doses of tamoxifen (5mg, gavage), only a small subset of cells was visible within the dermis and scattered in other connective tissues of the limb (Figure 2 E). Labelled Prrx1^enh+^ cells were found primarily within two specialized dermal niches: the hair follicle dermal papillae (dp) 84% (Figure 2 C) and in the perivascular space (Figure 2 D). Prrx1^enh+^ cells were also occasionally found in the dermal sheath of hair follicles and in papillary dermis. We confirmed in tissue sections that all Prrx1^enh+^ cells in the adult dermis were also PRRX1^+^. Prrx1^enh+^ cells made up just 0.22% ± 0.75 SD (n= 6) of the total PRRX1^+^ mesenchymal cells in the adult limb dermis. Likewise, dissociation of the dermis and FAC sorting revealed that approximately 0.48% ± 0.33 SD, of dermal cells were tdTOMATO^+^ (Figure 2 F). We also confirmed that the higher conversion dose required in young adult mice was not due to inaccessibility to tamoxifen, since administration either intraperitoneally or intravenously, yielded the same number of cells converted (data not shown). Furthermore, we dissociated dermal cells from *Prrx1enh-CreER;LSL-tdTomato* mice, to allow *in vitro* conversion of Prrx1^enh+^ cells and found no difference in the percentage of converted cells to that of intact dermis (data not shown). Thus, the Prrx1^enh+^ marks a broad embryonic population that reaches its peak during limb formation, but the number of cells that preserve the activity of the enhancer rapidly declines after birth. In contrast to the embryonic congruity of enhancer and protein, in adulthood, the Prrx1^enh+^ population becomes a small subset of the broad PRRX1 population.

**Figure 2.**
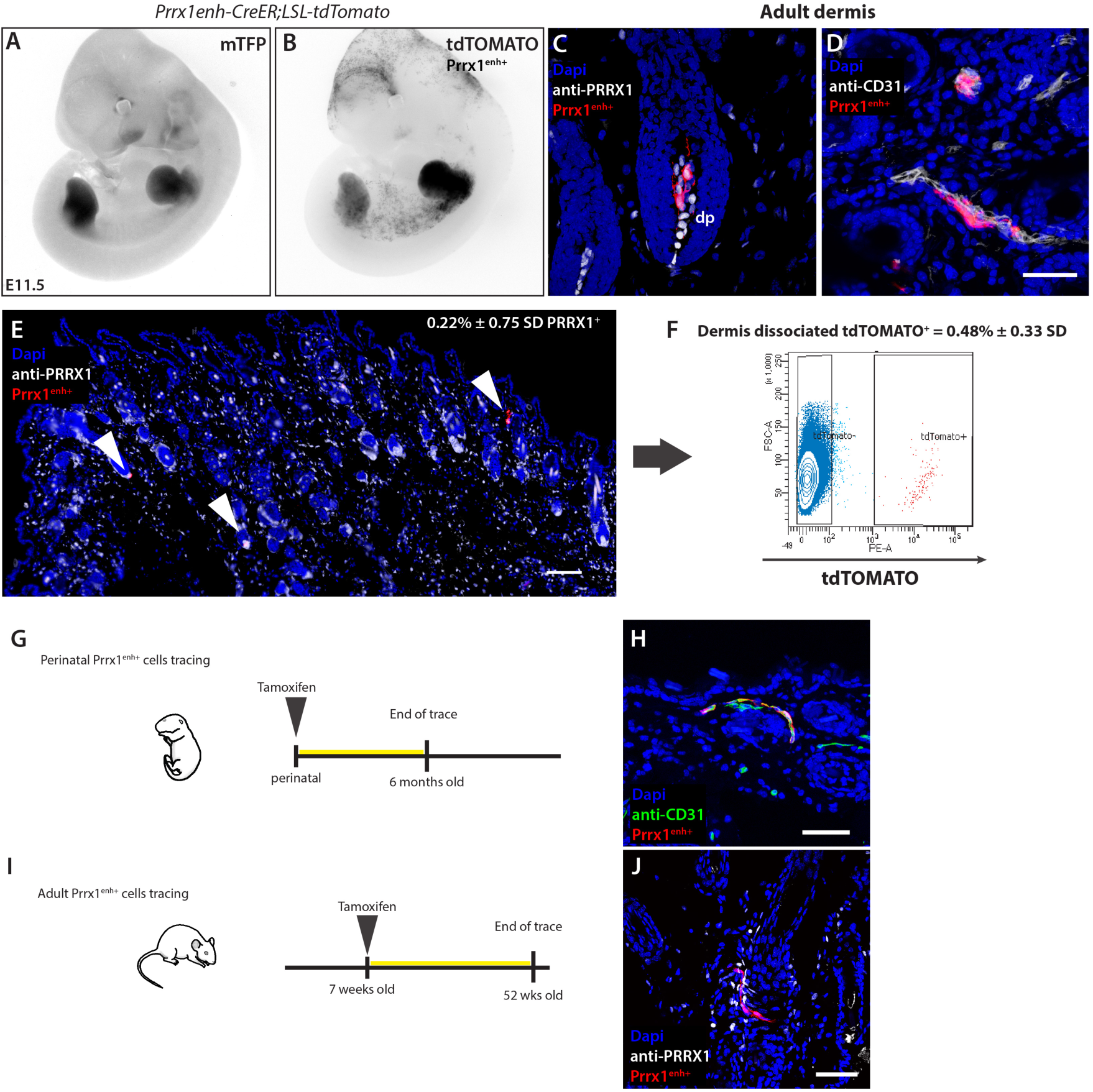
The *Prrx1* enhancer is active in limb bud mesenchyme progenitors and adult dermis. To lineage trace the fate of Prrx1^enh+^ cells, a transgenic *Prrx1 enhancer-CreER-T2A-mTFPnls* was crossed to a reporter line *Rosa-CAG-loxP-stop-loxP-TdTomato* (referred to as *Prrx1enh-CreER;LSL-tdTomato*). (A) Inverted greyscale images of embryos showing limb TFP expression at E11.5 under the *Prrx1* enhancer. (B) Upon tamoxifen administration (1mg oral gavage to mothers 24 hours prior to imaging), tdTOMATO^+^ (Prrx1^enh+^) is visible in limb buds. Additionally, Prrx1^enh+^ cells are present in a salt-and-pepper pattern in inter-limb flank, as well as cranial and craniofacial mesenchyme. (C, D) Prrx1^enh+^ cells in adult limb dermis. Cre recombinase activity labels cells primarily in dermal papilla (dp) and perivascular cells. Scale bar, 50 microns. (E) An overview of the sparsely-labeled cells within the adult limb dermis. Arrow heads mark Prrx1^enh+^ positive cells that make up 0.22%±0.75SD of PRRX^+^ cells. Scale bar, 200 microns. (F) Dermal dissociation of cells three weeks after tamoxifen administration and FAC sorting of tdTOMATO^+^ cells, shows that in adult skin only a small population of cells retain the activity of the PRRX1 enhancer. (G) Graphic representation of experimental set up to analyze the role of perinatal Prrx1^enh+^ cells in tissue homeostasis. (H) A representative tissue section of experiment (H) after 6 months. Scale bar, 50 microns. (I) Graphic representation of experimental set up to analyze the role of postnatal Prrx1^enh+^ in tissue homeostasis. (J) A representative micrograph of experiment (I) after 1 year. Scale bar, 50 microns.

An open question was if cells expressing the *Prrx1* enhancer at late embryonic and early postnatal periods would have a role in building and maintaining adult tissue. We administered tamoxifen perinatally and examined if Prrx1^enh+^ cell number would increase over time and in a clonal manner in specific dermal niches (Figure 3 G). We found that although the number of cells converted perinatally is higher than in adult, no clonal expansion was observed after 6 months (Figure 3 I). The same result was obtained when recombination was induced in 7 week-old mice and tissue was collected one year later (Figure 3 H, J). We next investigated if Prrx1^enh+^ adult cells might play a role in injury repair.

**Figure 3.**
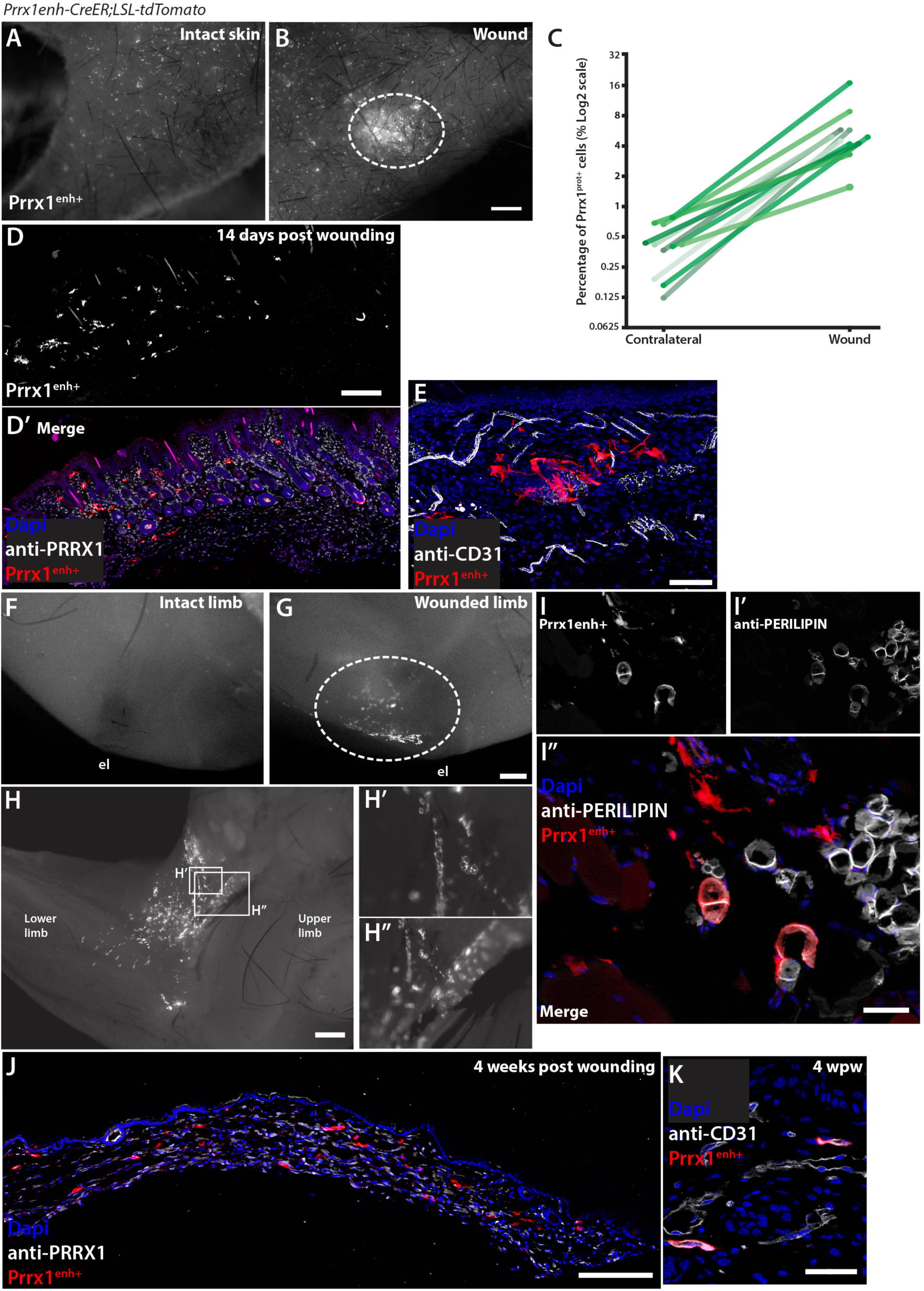
Prrx1^enh+^ cells amplify after injury in adult skin. (A) Stereoscope micrograph of skin from an intact limb. (B) Skin from wounded limb 21 days after wounding. Prrx1^enh+^ cells respond to 2 mm full thickness wounds in the limb, highlighted by dashed oval. Scale bar (A) and (B), 1mm. (C) Quantification of Prrx1^enh+^ cells in sections from paired samples of contralateral and wounded limb skin. The percentage of Prrx1^enh+^ from the total population of PRRX1^+^ cells/mm^2^ ± SD are plotted. Y axis is shown in log2 for optimal visualization of values below 1. (D, D’) A representative skin section of a wound 21 days after wounding. Scale bar, 200 microns. (E) Prrx1^enh+^ cell in the wound bed. Cells are not associated to blood vessels during the wound-resolving phase. Scale bar, 100 microns. (F) Fixed limb of mouse after skin collection in contralateral limb; (el) elbow. (G) Fixed limb of mouse after skin collection in wounded limb. Dashed oval marks the tissue under the wounded skin. Scale bar, 500 microns. (H, H’, H”) Wounded limb with inserts showing detail of Prrx1^enh+^. Scale bar, 1 mm. (I, I’, I”) Representative section of the limb subcutaneous tissue under the wound. In white is the adipocyte marker PERILIPIN. Scale bar, 50 microns. (J) At 4 weeks post wounding once most inflammation has resolved, Prrx1^enh+^ cells remain in the wound bed. Scale bar, 200 microns. (K) At 4 weeks post wounding Prrx1^enh+^ return to an association with blood vessels. Scale bar, 50 microns.

### Prrx1 enhancer cells are injury-responsive and amplify upon limb full-thickness skin wounding

We were curious if these limited Prrx1^enh+^ cells in the adult mouse retained embryonic-like properties that could contribute positively to repair and regeneration. Previously, the mouse *Prrx1* enhancer was shown to be active during wound healing and spike formation in Xenopus (Suzuki et al., 2007), but absent in wound healing of mouse back skin (Yokoyama et al., 2011), the predominant experimental paradigm for skin healing in the mouse. Although PRRX1 protein is present in dermal cells of back skin, we could detect no more than one or two Prrx1^enh+^ cells in multiple sections of at least 4mm expanding along the back (Supp. Fig. 1 A) (n=10 animals). The lack of enhancer positive cells in back skin was confirmed after tamoxifen administration in embryos, adult mice, or after injury.

Given that the *Prrx1* enhancer marks limb progenitors, and fibroblasts are known to carry unique positional information based on their location in different body parts, we decided to lineage trace Prrx1^enh+^ cells in a limb full-thickness skin injury model. We performed 2mm full thickness skin wounds in the upper limbs of *Prrx1enh-CreER;LSL-tdTomato* (Supp. Fig. 1 B). Mice were wounded three weeks after the last administration of tamoxifen to prevent recombination during wound healing. In contrast to back skin wounds, it was not possible to splint limb skin wounds. A semi-occlusive bandage (Tegaderm 3M) was applied for the first 48-60 hours to prevent infection. This bandage only mildly impedes wound contraction that is characteristic of skin wounds in rodents (Supp. Fig. 1 C). Fourteen days post wounding (14 dpw), animals were sacrificed and the wounded and contralateral limb were compared for the percentage of Prrx1^enh+^ (tdTOMATO^+^) cells. At low magnification there was already an obvious amplification of Prrx1^enh+^ cells within the wound bed compared with the adjacent wound margin and contralateral limb (Figure 3 A, B). Within 1mm^2^ cross sections of the wound (Figure 3 D, D’), despite an increased density of mesenchymal PRRX1^+^ cells, the percentage of labeled Prrx1^enh+^ cells were on average 16.46-fold increased (±12.58 SD, *p*=0.0015) over the uninjured contralateral dermis (Figure 3 C). This represents an increase of Prrx1^enh+^ from 0.37 % ±0.2 SD to 5.76 % ± 3.8 SD of the total PRRX1^+^ cells. During the injury response, Prrx1^enh+^ cells were not necessarily associated to blood vessels (Figure 3 E), but primarily within the upper or papillary dermis. These results were confirmed in three independent experiments.

An important question to address was if other subpopulations of dermal cells would also amplify in response to injury in a similar manner as Prrx1^enh+^ cells. To address this, we used a transgenic mouse expressing inducible CreER under the control of a 17-kb fragment of the *Col1a2* upstream enhancer crossed with a *Rosa-CAG-loxP-stop-loxP-TdTomato* mice (referred to as *Col1a2enh-CreER;LSL-tdTomato*). This transgenic mouse has been used previously as a reporter of wound healing in dorsal wounds (Higashiyama et al., 2011; Rajkumar et al., 2006). We administered tamoxifen 3 weeks before wounding and quantified tdTOMATO^+^ cells in contralateral and wound tissue (Supp. Fig 2 A, B). In uninjured tissue Col1a2^enh+^ cells labeled dermal fibroblasts as well as portions of surface and interfollicular epithelia. No difference was found in the percentage of Col1a2^enh+^ cells among the total of mesenchymal PRRX1^+^ cells, in injured and intact skin (Supp. Fig 2 D). In contrast to Prrx1^enh+^ cells, we found no enrichment of positive dermal cells within the wound bed or in the subcutaneous space under the wounds (Supp. Fig 2 G, H). In fact, there was a small depletion of Col1a2^enh+^/PRRX^+^ within the wound 21 days after injury. This suggests that the dermal cells labeled by Col1a2^enh+^ have a muted response to injury and may be slightly displaced by other subpopulations including Prrx1^enh+^ cells.

### In response to injury, dermal *Prrx1*-enhancer cells contribute to tissues beyond dermal compartments

When collecting skin for tissue processing, we observed a concentration of Prrx1^enh+^ positive cells in sub cutaneous connective tissue surrounding the muscle immediately under the wounded skin (Figure 3 F, G, H-H”). We found that Prrx1^enh+^ cells were located within fascia, loose connective tissue and adipose tissue. We confirmed the contribution to adipose tissue by co-staining Prrx1^enh+^ with a PERILIPIN antibody (Figure 3 I-I”). In our experimental layout, tamoxifen is administered 5 weeks prior to tissue collection and recombination labels only a dispersed, small subset from the broad PRRX1^+^ population in the unwounded or contralateral limb. This suggests that upon injury, labeled, dermal-associated cells migrate from the dermis and take up residence in subcutaneous tissue where they contribute to diverse tissue types such as adipose tissue. After 4 weeks, when the 2 mm wound is resolved, and cellular content is decreased, Prrx1^enh+^ cells remain in the wound bed (Figure 3 J) where they re-acquire an association with blood vessels (Figure 3 K). The number of Prrx1^enh+^ cells in the wound after 4 weeks remains elevated compared to contralateral uninjured skin by 11.48-fold increase (± 4.1 SD, n=6). The vast Prrx1^enh+^ progeny in the wound suggests that cells that have an active enhancer under homeostatic conditions readily migrate into the wound bed and proliferate relative to other PRRX1^+^ cells.

*Prrx1* has been shown to be a marker necessary for maintaining stemness in adult hippocampal tissue (Shimozaki et al., 2013). We hypothesized that cells expressing *Prrx1* would contribute to wound repair by either providing daughter cells that would differentiate and directly contribute to structuring the wound, or by supplying factors that modulate healing.

As a marker of regenerative progenitors, *Prrx1* upregulation has been viewed as a stereotypical step in the formation of the proliferative blastema during salamander limb regeneration (Satoh et al., 2007). Recent lineage tracing in the axolotl showed that Prrx1^enh+^ cells are multipotent cells that can regenerate skeleton and soft connective tissue (Gerber et al., 2018). The molecular mechanism underlying the enrichment of Prrx1^enh+^ cells within mouse skin wounds is unknown, but live imaging of connective tissue during axolotl regeneration uncovered a pivotal role of cell migration in the process of blastema cell creation (Currie et al., 2016). It is tempting to think that activation of the *Prrx1* enhancer could relate to an increased readiness or propensity to migrate, which could relate to the role of *Prrx1* in epithelial to mesenchymal transitions (EMT) (Ocaña et al., 2012). In this study, we show that the protein expression of *Prrx1* differs from the activity of the 2.4kb upstream enhancer, and the results presented here argue for a subpopulation of enhancer versus PRRX1^+^ cells.

The heterogenous responses to healing by different fibroblastic populations is inarguably complex, and at least three sources could account for some aspects: 1) embryonic origin, 2) location in the body and 3) the microenvironment. One illustrating example is the different response to regeneration from fibroblasts in the P3 phalanx versus fibroblasts in the P2 phalanx (Wu et al., 2013).

In this work we investigated the response of *Prrx1* enhancer-expressing cells to injury and homeostasis. Because this enhancer is limb specific, we focused on full-thickness skin wounds in the limb. The majority of wound healing assays in murine models are based on back skin wounding due to the inaccessibility during grooming, the ability to make large wounds surpassing 5mm in diameter, and the possibility to splint (to mimic wound healing in humans where there is no contraction). The embryonic source of dorsal cells is diverse and could encompass cells from neural crest, pre-somitic mesoderm, and lateral plate mesoderm. In contrast, connective tissue of the limbs derives primarily from a single embryonic source of lateral plate mesoderm. In the future it will be interesting to assess if limb dermal cells display less heterogeneity compared to the dermal compartment of other regions that derive from diverse embryonic origins. In addition to being of a single embryonic source, experimental paradigms of wound healing in limbs have an important and clinical relevance for diabetic and trauma-related injuries.

One of the most surprising results was the wide contribution of Prrx1^enh+^ cells to subcutaneous tissues under the wound bed. We administered Tamoxifen three weeks prior to wounding and never observed an enrichment of subcutaneous cells except in the context of wounding and did not observe a similar phenotype when labeling Col1a2^enh+^ fibroblasts. This suggests to us that Prrx1^enh+^ cells from the dermis emigrate into foreign subcutaneous tissues and contribute to adipocytes, fascia, and other structures. This resident tissue plasticity may be a conserved feature of Prrx1^enh+^ cells, since axolotl Prrx1^enh+^ cells are able to contribute to new segment formation in contrast to other mesenchymal cell populations (Gerber et al., 2018).

While Prrx1^enh+^ cells are able to expand and contribute to healing tissue, they do not seem to play a role in normal tissue growth and homeostasis as is the case for most epithelial stem cell pools. By labeling at several prenatal and early postnatal timepoints, we were unable to observe long term expansion of Prrx1^enh+^ clones that might arise from a tissue resident stem cell. Instead, Prrx1^enh+^ cells remained relatively quiescent during post-embryonic growth and restricted to the perivascular and dermal papilla niches. It was only post-injury that cells amplified during wound healing migration and proliferation. Our work suggests that Prrx1^enh+^ cells represent a specialized pool of injury-responsive cells. Recent work has highlighted similar populations of cells that are specifically tuned for tissue repair but not tissue maintenance and growth (Llorens-Bobadilla et al., 2015; Lopez-Baez, 2018; Marecic et al., 2015). It will be interesting in the future to understand what molecular factors drive the *Prrx1* enhancer during injury and to what degree they overlap with regulators of embryonic *Prrx1* enhancer activity. Based on the differences in *Prrx1* enhancer activity and PRRX1 protein expression, we hypothesize that there may be distinct molecular differences that differentiate enhancer-positive cells from the broader PRRX1 population which manifests in large differences in cell behavior during wound repair.

Overall, our results highlight a unique injury-responsive cell population within adult tissue. As impressive as the response of Prrx1^enh+^ cells to injury, their amplified numbers still make up only a small fraction of the overall population of wound fibroblasts. Future work aimed at understanding the molecular signals that retain and specify Prxx1^enh+^ cells will be key to tipping the balance from scar formation to regenerative tissue replacement.

## Materials and Methods

### Transgenic mouse lines

A construct was created containing the 2.4kb mouse *Prrx1* enhancer (REF) followed by the b-globin intron and nuclear-localized teal fluorescent protein-1 (mTFP1-nls), a T2A self-cleaving peptide, and CreERT2. The construct was linearized by digestion by KpnI and injected into the pro-nucleus of Bl6/J mouse oocytes (MPI-CBG Transgenic Core Facility). Six founder animals were generated and F1 progeny were screened for mTFP expression during embryonic limb development (E10.5-E12.5) and the ability to induce recombination of LoxP reporter lines only after administration of tamoxifen. From the initial founder animals, one line was further characterized and used for all subsequent experiments. Further lines used in this study: B6;129S6-Gt(Rosa)26Sor^*tm9(CAG-tdTomato)Hze*^/J (Jackson Laboratory), B6.129S4-Pdgfra^*tm11(EGFP)Sor*^/J (Jackson Laboratory), B6.Cg-Tg(Col1a 2-cre/ERT,-ALPP)7Cpd/J (Jackson Laboratory). All mouse lines were bred in MPI-CBG, CRTD and IMP facilities. All procedures performed in animals adhered to local ethics committee guidelines.

### Immunohistochemistry

Full thickness skin was dissected from mouse upper forelimb or lower back and attached to filter paper before submersing in 4%PFA in phosphate buffer. Tissue was fixed overnight at 4°C and then washed in PBS and serial overnight washes in 10% and 30% sucrose in PBS before being embedded in OCT and cryo-sectioned in longitudinal sections of 12 µm. For mouse limbs from E18.5 to 52 weeks of age, limbs were harvested and fixed overnight in PFA, washed in PBS, and incubated with 400 µM EDTA in PBS for approximately one week before washing in sucrose, embedding and cryo-sectioning. Standard immunohistochemistry techniques were used for staining. Primary antibodies: PRRX1 (MPI-CBG Antibody facility) used at 1:200, SOX9 (R&D systems AF3075) used at 1:200, PERILIPIN (Sigma-Aldrich P1873) used at 1:200, CD31 (BD Pharmingen 557355) used 1:100.

### Wounding

Animals were anesthetized by intraperitoneal injection of a mixture of Ketamine (100 mg/kg) and Xylazin (10 mg/kg). Skin was shaved, chemically defoliated, and wiped with 70% ethanol. To create a 2 mm wound in the forelimb, we pulled the skin from the posterior part of the limb (skin in the forelimb is not attached to the muscle-skeletal core) and at the crease, perforated with the half of a 2 mm punch biopsy. Wounds were immediately disinfected with iodine solution and wrapped with Tegaderm (3M) and bandage. Animals were given Carprofen at 4mg/kg and monitored during anesthetic recovery. Wounded animals were monitored daily for signs of infection and any self-removal of bandages. If animals had not removed forelimb bandages and Tegaderm by two to three days post wounding, the dressing was manually removed to synchronize the kinetics of wound resolution. The wound in the posterior area of the limb ensured that the mice were not able to reach it and affect the healing process. We did not observed infections or any complication due to the wound. Animals were sacrifice at either 5, 14, 21, or 28 days post wounding and skin was processed as described above.

### Tissue dissociation and FACS

Animals were sacrificed and upper arm skin was shaved and cut as full thickness skin. Adipose depots were manually removed, and the skin was washed once with 70% ethanol and three times with cold, sterile PBS. Skin was incubated in 10mg/ml elastase in DMEM at 37C° for 20 minutes. Dermal tissue was manually removed and separated from epithelial tissue. Dermal tissue pieces were then placed in 0.35mg/ml Liberase TM in PBS with 10 Units/µl DNase for 37C° for 30 minutes and then manually dissociated by mechanical disruption with forceps to achieve a single cell suspension. Cell suspensions were filtered through a 50µm mesh and diluted in 10% serum containing DMEM. Cell suspensions were centrifuged at 900rcf and resuspended in serum-containing DMEM. Both filtered cells and unfiltered tissue pieces were plated on gelatin coated-tissue culture plates. 24 hours post-dissociation both filtered cells and unfiltered tissue were visually assessed for tdTOMATO expression. 24-48 hours after dissociation, cells were trypsinized, washed, and resuspended in FACs buffer containing 1x PBS (Ca/Mg free), 2mM EDTA, 25mM Hepes (pH 7.2), 1% BSA, and pen/strep. Non-transgenic controls skin was used to determine gating for tdTOMATO signal.

### Imaging

Images were acquired with a 20x objective Apotome 2 (Zeiss) or inverted laser scanning confocal 780 (Zeiss) at 20 or 40x magnification. Imaging was performed on instruments of the Light Microscopy Facility, a core facility of CRTD at Technische Universität Dresden. Images were analyzed using Fiji and Photoshop, and alterations to brightness or contrast were applied equally to the entire micrograph for visualization purposes only. Cell quantification was performed in two sections per wound, of the mid area, of each animal. A region of interest measuring 1mm^2^ was cropped in Fiji for quantification. Wound sections were paired with their contralateral control to determine the fold increase and investigator was blinded to the sample grouping. Statistical analyses were performed using the Graphpad Prism 7.0 software.

## Author Contributions

JDC and TSG conceived and executed experiments, analyzed data, and wrote the manuscript. LG, MM and MS contributed experimental data and support. PM contributed the *Prrx1* enhancer construct and antibody. EMT and TSG secured funding for the project.

## Acknowledgements

This work was supported by the animal facilities of the MPI-CBG, CRTD and IMP with institutional funding to EMT and TSG. EMT is supported by an ERC Advanced Grant. JDC acknowledges support from an EMBO Long Term Fellowship, Alexander Von Humboldt Fellowship, and MPI
-CBG Fellowship. This work was supported by the Light Microscopy Facility, a core facility of BIOTEC/CRTD at Technische Universität Dresden. The authors claim no conflict of interest.

