## Supplemental figures for "The *Prrx1* limb enhancer marks an adult population of injury-responsive dermal fibroblasts"

### SUPPEMENTARY DATA

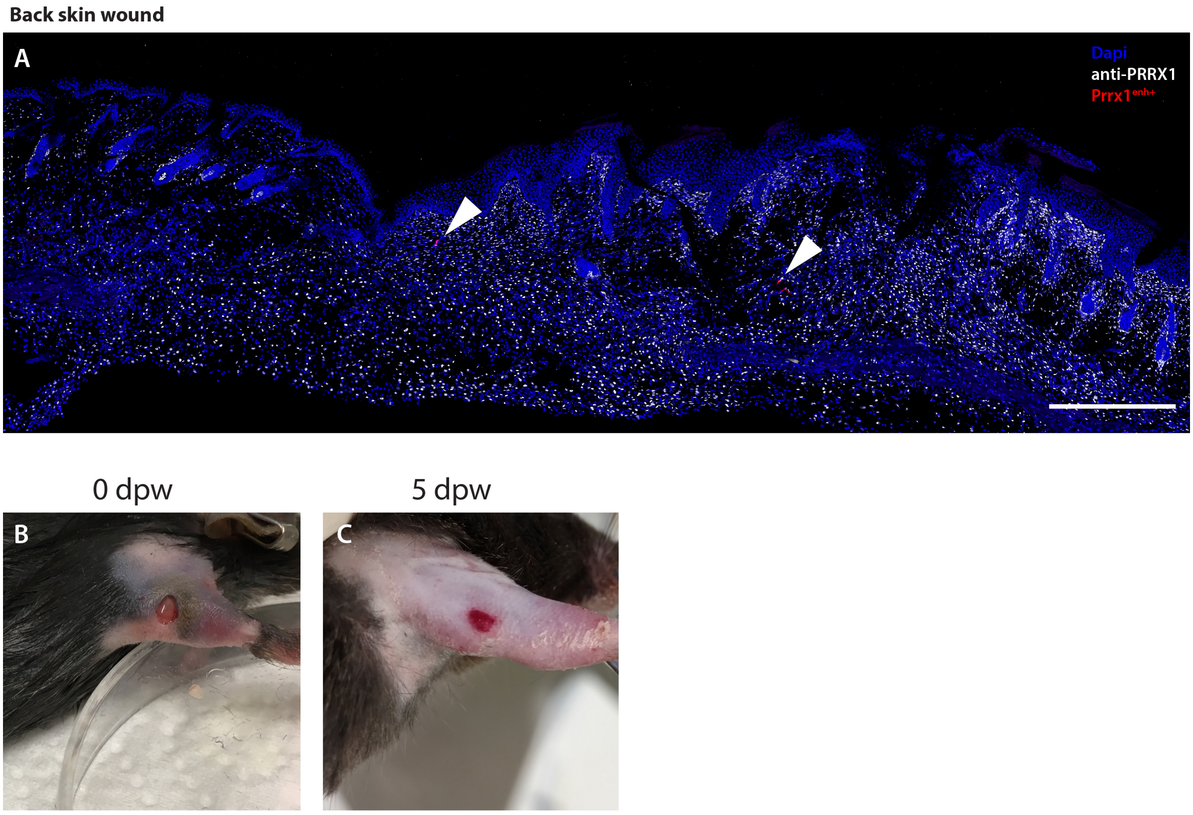


**Figure S1.**

(A) 2 mm full-thickness wounds in the dorsal trunk of *Prrx1enh-CreER;LSL-tdTomato* mice produce 0-10 labeled cells in a cubic centimeter of wound tissue. Scale bar, 500 microns.

(B) A fresh 2 mm full-thickness wound in the posterior skin of the upper limb. The position of the wound does not enable splinting of the wound, therefore, we used semi-occlusive dressing to protect the wound from infection and delay the contraction of the wound.

(C) After 5 days post wounding the semi-occlusive dressing is removed as healing proceeds.

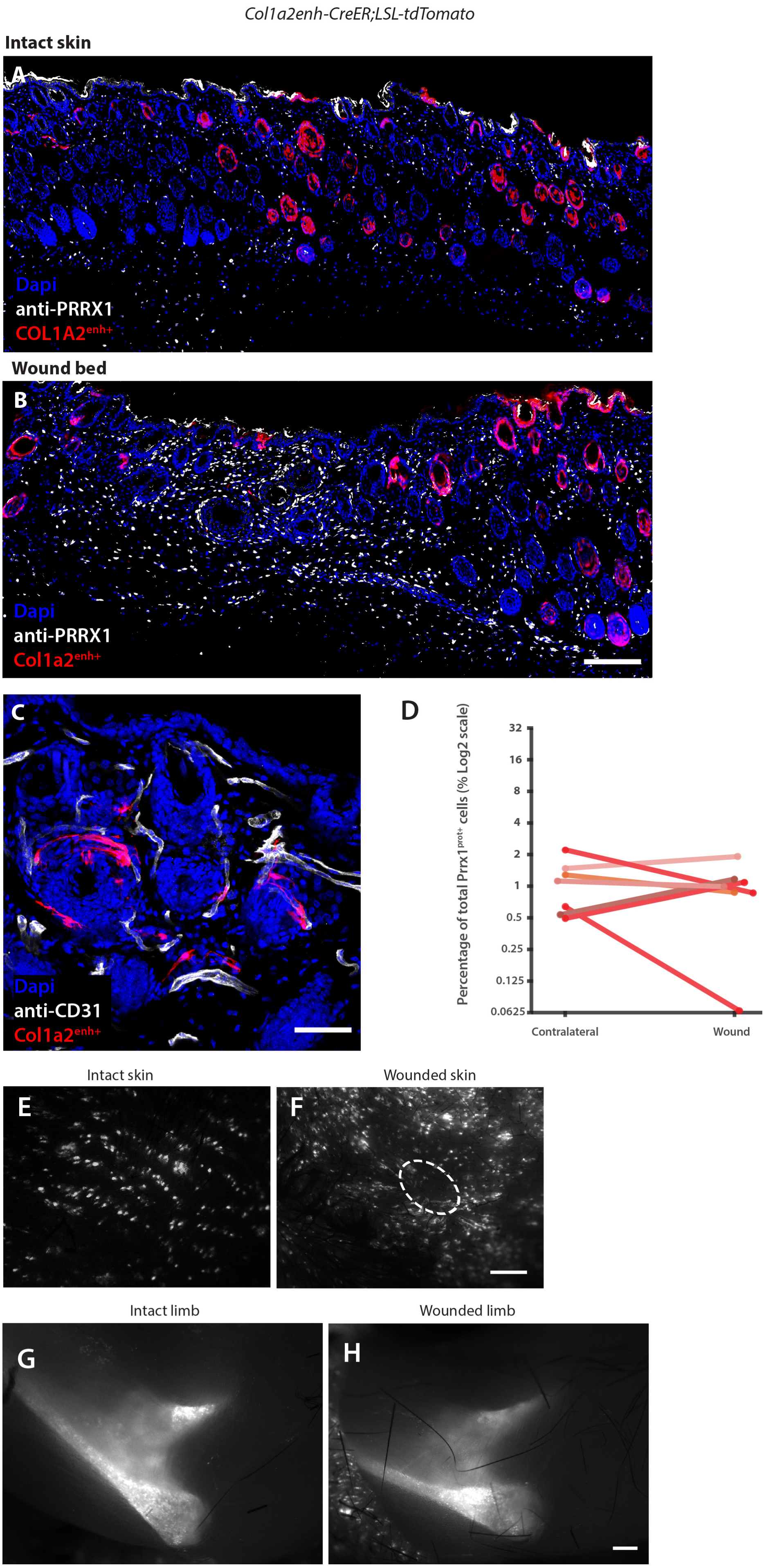

Col1a2enh-CreER;LSL-tdTomato
Intact skin
A
Dapi
anti-PRRX1
COL1A2enh+
Wound bed
B
Dapi
anti-PRRX1
Col1a2enh+
C
Dapi
anti-CD31
Col1a2enh+
D
Percentage of total Prrx1prot+ cells (% Log2 scale)
32
16
8
4
2
1
0.5
0.25
0.125
0.0625
Contralateral
Wound
Intact skin
Wounded skin
E
F
Intact limb
Wounded limb
G
H

**Figure S2.**

(A) Intact skin of *Col1a2enh-CreER;LSL-tdTomato* mice 3 weeks after administration of tamoxifen. The majority of the labelled cells are in epidermal layers with some dermal cells visible. Scale bar, 200 microns.

(B) A 2 mm full thickness wounds in the posterior skin of the upper limb of *Col1a2enh-CreER;LSL-tdTomato.* $Col1a2^{enh+}$ cells are mostly absent in the wound bed. Scale bar, 200 microns.

(C) $Col1a2^{enh+}$ cells are not associated to blood vessels. Scale bar, 50 microns.

(D) Quantification of $Col1a2^{enh+}$ cells in sections from paired samples of contralateral and wounded limb skin. The percentage of $Col1a2^{enh+}$ from the total $PRRX1^{+}$ of cells/$mm^2$ ± SD are plotted. Y axis is shown in log2 for optimal visualization of values below 1.

(E) Stereoscope micrograph of skin from an intact limb.

(F) Skin from wounded limb 21 days after wounding. $Col1a2^{enh+}$ cells are mostly absent in 2 mm full thickness wounds in the limb, highlighted by dashed oval. Scale bar, 1 mm.

(G) Fixed limb of mouse after skin collection in contralateral limb.

(H) Fixed limb of mouse after skin collection in wounded limb. Positively labeled cells in the bone and skin and absent in superficial subcutaneous tissue under the wounded dermis. Scale bar, 500 microns.
